# The cortical contribution to the speech-FFR is not modulated by visual information

**DOI:** 10.64898/2026.01.26.701703

**Authors:** Jasmin Riegel, Alina Schüller, Alexander Wißmann, Steffen Zeiler, Dorothea Kolossa, Tobias Reichenbach

## Abstract

Seeing a speaker’s face can significantly aid understanding, particularly in challenging acoustic environments. An early neural response implicated in audiovisual speech processing is the frequency-following response (speech-FFR), which occurs at the fundamental frequency of the speech signal. This response arises from both subcortical areas and the auditory cortex. Previous studies have shown that subcortical responses are reduced when bimodal stimulation includes visual input from the talker’s face. Here, we examined the cortical contribution to the speech-FFR and its potential modulation by visual information. We recorded MEG responses to four types of audiovisual signals: a still image, an artificially generated avatar, a degraded video, and a natural video. The audio stimuli were presented in a substantial level of background noise to make behavioral audiovisual effects stand out. Speech-in-noise comprehension increased significantly from the audio-only condition to the avatar and the degraded video, and further to the natural video. Moreover, we found that all types of audiovisual stimuli yielded robust speech-FFRs in the auditory cortex at an early latency of around 30 ms. However, the magnitude of this neural response was neither enhanced nor attenuated by the videos, nor could the cortical contribution of the speech-FFR explain a significant portion of the variance in the behavioral comprehension scores. Our results suggest that visual modulation of the speech-FFR in the auditory cortex is, if existent, too small to be measurable in scenarios where speech occurs in considerable background noise.

## Introduction

Understanding speech in acoustically challenging environments can be difficult, not only for individuals with normal hearing but especially for those with hearing impairments [Miles et al., 2022]. Speech com-prehension can be substantially enhanced when listeners see the speaker, since visual information, such as lip movements and facial cues, supports auditory speech processing [Thézé et al., 2020, Nirme et al., 2020]. The improvement in speech comprehension is most notable in the presence of background noise, a principle known as inverse effectiveness [Sumby and Pollack, 1954, Grant and Seitz, 2000, Ross et al., 2007].

Numerous studies have investigated the neural mechanisms underlying audiovisual speech integration. Most of this work has focused on event-related potentials [Van Wassenhove et al., 2005, Pilling, 2009] or on cortical tracking of the low-frequency fluctuations in the speech envelope [Crosse et al., 2015, 2016, Varano et al., 2025]. It identified, amongst others, a role of the delta frequency band in linking auditory and visual information on the word level [Varano et al., 2025, Riegel et al., 2026].

However, neural responses to speech also occur in the much higher gamma frequency range. In particular, neural activity can phase-lock to the fundamental frequency of speech, as well as to a lesser extent, to its higher harmonics. This frequency-following response (speech-FFR) has initially been assessed in response to single repeated syllables [Russo et al., 2004, Skoe and Kraus, 2010]. Recent studies have shown how, through statistical modeling, the speech-FFR can also be detected in response to continuous, non-repetitive speech in which the fundamental frequency varies over time [Forte et al., 2017, Etard et al., 2019].

The speech-FFR, as measured with electroencephalography (EEG), has largely subcortical origins, including in particular the inferior colliculus. The speech-FFR has therefore been conceptualized as reflecting predominantly feedforward, audio-stimulus-driven encoding in the auditory brainstem and midbrain [Cof-fey et al., 2019]. However, there have also been reports of modulatory influences beyond purely feedforward processing. In particular, a series of studies suggested that long-term musical training can enhance the fidelity of subcortical FFRs to speech and non-speech sounds [Musaccia et al., 2007, Musacchia et al., 2008]. This interpretation has recently been challenged by a large-scale, preregistered study that failed to find evidence for such effects of musical training [Whiteford et al., 2025].

In contrast to experience-related effects such as musical training, the extent to which the subcortical speech-FFR is modulated by top-down attention remains debated [Lehmann and Schönwiesner, 2014, Forte et al., 2017, Stoll et al., 2025]. Of particular interest for audiovisual speech perception, audiovisual stimuli have been found to suppress the magnitude of the speech-FFR [Musacchia et al., 2006].

Coffey et al. [2016] used magnetoencephalography (MEG) to show that the speech-FFR elicited by repeated syllables contains a reliable cortical contribution, in addition to the responses from the brainstem and the midbrain. Kulasingham et al. [2020] added to this finding by showing that the cortical contribution to the speech-FFR could also be identified when elicited by continuous speech. Since then, factors that potentially impact the subcortical speech-FFR have increasingly been investigated regarding their effect on the cortical contribution. In particular, we and others recently demonstrated sizable and robust attentional modulation of the speech-FFR in the auditory cortex [Schüller et al., 2023, Commuri et al., 2023]. On the other hand, we were unable to find an effect of musical training on this response [Riegel et al., 2024].

In the following study, we investigated whether the previously reported audiovisual suppression of the sub-cortical contribution to the speech-FFR extends to the cortical contribution. To investigate this, we measured the neural response to continuous audiovisual speech in 32 participants using MEG. Stimuli were presented in four conditions: a still image of the speaker, an avatar, a degraded version of the speaker, and the natural video.

We included an artificially generated avatar in the experiment because visual access to a speaker’s face is not always available, for example, in public settings such as train station announcements. In such cases, avatars may offer a promising alternative by providing visual speech cues to support comprehension. Previous be-havioral studies have shown that avatars can partially improve speech understanding in noisy environments, although typically to a lesser extent than natural visual speech [Varano et al., 2022, Nirme et al., 2020, Shan et al., 2022].

The degraded video was included to determine the putative effect of simplified visual signals on the cortical speech-FFR, that is, of visual signals that contain sufficient information to enhance speech-in-noise com-prehension, although to a lesser extent than natural videos. To maximize the potential audiovisual benefit – and, consequently, the expected effect size in the cortical speech-FFR – we followed the principle of inverse effectiveness and embedded the audiovisual speech stimuli in four-talker babble noise.

## Materials and methods

We used the data from a previous MEG study on audiovisual speech perception [Riegel et al., 2026]. Thirty-two participants listened to a single speaker uttering sentences in background noise. After each sentence, the participants repeated what they understood. The audiovisual speech material was presented in four different visual conditions: (1) a still image of the speaker, (2) a synthetically generated avatar, (3) a degraded video, and (4) the natural video. While our previous study focused on tracking of low-frequency amplitude fluctuations of speech in the auditory cortex, in the current investigation, we determined the higher frequency tracking of the presented speech in the auditory cortex, namely the speech-FFR.

### Audiovisual stimuli

The speech material was based on recordings of a native German speaker uttering short sentences from the Oldenburger Satztest, a standardized matrix-sentence corpus designed to generate grammatically correct and phonetically balanced but semantically unpredictable sentences [Wagener et al., 1999]. Each sentence follows the same syntactic structure - name, verb, number, adjective, and object. The lexical items are randomly drawn from sets of ten alternatives per position, resulting in one million possible unique sentences.

Recordings were conducted at *29.97* fps with accompanying audio sampled at *44.1* kHz. We collected 600 unique sentences and screened them to ensure consistent pronunciation and the absence of background noise. After this quality control step, 571 sentences remained and served as the basis for generating the four stimulus types.

To create the degraded audiovisual condition, we processed the original videos using the *edgedetect* function from FFmpeg, which emphasizes prominent contours while suppressing surface detail.

The avatar-based condition was generated by combining a still image of the speaker with the corresponding audio file using software provided by D-iD [D-ID, 2025]. The image was animated by applying a convolutional neural network to encode the speaker’s facial features and a generative adversarial model to animate the face in synchrony with the speech signal.

All sentences were presented in background noise to increase the potential effects of audiovisual integration [Ma et al., 2009]. We employed four-talker babble noise that began shortly before sentence onset. We determined the signal-to-noise ratio during a pilot study, aiming for approximately 50% comprehension in the audio-only condition. This procedure resulted in an SNR of approximately *-4* dB and an overall sound pressure level of 68 db(A). We used this level uniformly for all participants.

### Experimental procedure

Thirty-three healthy, right-handed adults (17 female, 16 male; age range 19–31 years) were recruited for the experiment. Each participant completed two sessions of up to 60 minutes, scheduled one to three weeks apart. One female did not finish both sessions, leaving data from 32 participants for analysis.

To familiarize the participants with the restricted vocabulary and thereby minimize later learning effects, the first MEG session started with 80 audio-only sentences with adaptive background noise, estimating each participant’s individual SRT_50_. Following the training, participants were exposed to the different types of audiovisual stimuli. The experiment was designed in blocks of sentences, with each block containing three randomly selected sentences from each condition. The order of all stimuli within them was fully randomized. Sentences were not grouped by condition. For every stimulus type, 60 sentences were presented. Individual sentences lasted approximately three seconds, yielding roughly three minutes of material per condition.

After each sentence, participants repeated all words they recognized while viewing the OLSA response matrix.

### Data acquisition and preprocessing

MEG recordings were obtained with participants lying supine and with eyes open using a 248-channel whole-head system (4D-Neuroimaging). The data was recorded at a sampling rate of 1,017.25 Hz in a magnetically shielded room. During acquisition, an online analog band-pass filter from 1 to 200 Hz was applied. The system further incorporated a calibrated linear combination of 23 reference sensors from 4D Neuroimaging (San Diego, CA, USA), enabling effective suppression of environmental magnetic interference. Before each session, we digitized the head shape along with five fiducial points using a Polhemus tracking system.

Subsequent offline preprocessing was carried out in MNE-Python [Gramfort et al., 2014]. We removed power-line artifacts with a 50 Hz notch filter (zero-phase FIR, transition bandwidth 0.5 Hz) and then resampled the data to 1,000 Hz.

The auditory stimuli were presented via a custom setup designed by Schilling et al. [2020]. To ensure precise timing, the audio signal was simultaneously routed to an MEG channel, and its alignment with the presented sound was obtained by cross-correlation, yielding temporal precision on the order of 1 ms. Visual material was shown on a screen placed above the participant. A constant delay between audio and video of a presented sentence was introduced by the audiovisual presentation setup. This delay was measured and corrected in the stimuli following the procedure described in [Varano et al., 2023].

### Neural Source Localization

We reconstructed neural sources using MNE-Python [Gramfort et al., 2014] together with the FreeSurfer “fsaverage” brain [Fischl, 2012]. To align this template to each participant, we co-registered it to the digitized head shape and fiducial points by applying rotations, translations, and uniform scaling based on the recorded head position data.

From the co-registered anatomy, we generated a volumetric grid with 5 mm spacing. Cortical regions of interest were then defined according to the Aparc atlas and encompassed bilateral temporal areas known to contribute to audiovisual speech processing [Van Atteveldt et al., 2010, Macaluso et al., 2004]. These ROIs included the inferior, middle, and superior temporal gyri, the transverse temporal gyri, the banks of the superior temporal sulci, as well as the fusiform, entorhinal, parahippocampal, and temporal pole regions. In total, this produced 862 source voxels (432 in the left hemisphere). Forward models were computed using a boundary element model (BEM).

To estimate source activity, we applied a linearly constrained minimum variance (LCMV) beamformer [Bourgeois and Minker, 2009]. The spatial filters were computed using the empirical covariance of the MEG data and the noise covariance derived from empty-room recordings. Filters were normalized to achieve unit noise gain, and their orientation was constrained to surface-normal directions.

### Behavioral Analysis

After each presented audiovisual stimulus, participants repeated every word they understood. For each stimulus type, the averaged speech comprehension of each participant was calculated.

### Linear forward models to determine neural speech tracking

The speech-FFRs were computed by estimating the neural response from two speech features, the fundamental waveform and the envelope modulation of the higher harmonics. The fundamental frequency was computed from the speech signal using the probabilistic YIN-algorithm [Mauch and Dixon, 2014]. The range of the estimated *f*_0_ (70–120 Hz) was used to define the passband edge frequencies of a bandpass filter applied to the speech stimuli in order to extract the fundamental waveform. The envelope modulation of the higher harmonics, first described by [Hertrich et al., 2012], was obtained by first applying a model of the auditory periphery and subsequently filtering the resulting signal in the same frequency range as the fundamental frequency [Chi et al., 2005]. In detail, the model begins by applying a set of constant-Q bandpass filters, which are designed to provide frequency-dependent resolution across the auditory spectrum. The filter outputs are subsequently processed using nonlinear compression and multiscale derivative operations, leading to enhanced spectral selectivity. Next, 86 frequency channels corresponding to harmonics above 300 *Hz* were selected. Within each channel, the amplitude envelope was extracted and bandpass filtered around the fundamental frequency *f*_0_ (70–120 Hz). Finally, these filtered envelopes were averaged across all channels to produce a single high-frequency envelope modulation feature.

The temporal response functions were derived using the stimuli brain mapping toolbox sPyEEG [Guilleminot et al., 2026]. The two speech features were then used as two predictors *x_i_*(*t*), with *i* ∈ {1, 2}, to estimate the MEG response *y*^(^*^j^*^)^(*t*) at voxel *j* and time *t* through a linear model with temporal delays:

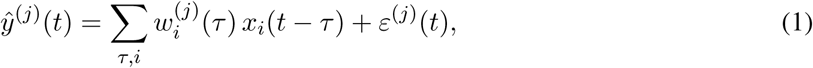

where 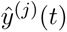 denotes the predicted MEG response, *x_i_*(*t* − *τ* ) is the *i*th stimulus feature at time lag *τ* , the weights 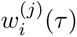 are the TRFs for the *i*th predictor variable and the neural activity at voxel *j* at time lag *τ* , and *ε*^(*j*)^(*t*) is the residual error term.

For training the model weights, ridge regression was applied with a normalized regularization parameter, such that the cost function reads:

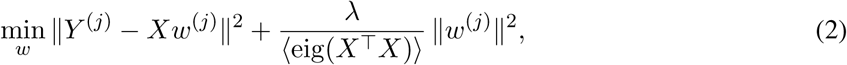

where *X* is the matrix of predictor variables at different time lags, *Y* ^(*j*)^ the source-reconstructed measure-ment data at voxel *j*, *λ* the defined regularization parameter, and ⟨eig(*X*^⊤^*X*)⟩ the mean eigenvalue of *X*^⊤^*X* used to normalize the regularization strength.

A range of regularization parameters was applied, *λ* = [10^−3^*, …,* 10^3^]. The regularization parameter yield-ing the highest reconstruction score was estimated using five-fold cross-validation and Pearson correlation for each of the four audiovisual conditions. This yielded *λ* = 10. A slightly higher value of *λ* = 100 was chosen as the optimal parameter to additionally ensure the avoidance of overfitting. Before computing the linear forward model, both stimulus and MEG data were z-scored and concatenated for each participant and condition.

The TRFs were calculated for each of the 862 voxels between −50 ms and 170 ms. The absolute values of the resulting TRFs were averaged across voxels to obtain an individual TRF for each participant, condition, and predictor variable. For the population-level TRFs, the average across participants was computed. The time lag of the peak in this response was identified. For the statistical analysis, the peaks in the TRFs of the individual participants were determined within a ±50 ms range around the latency of the peak in the population average.

### Statistical Analysis

We first tested whether the TRFs at the population level differed significantly from noise. We therefore generated a noise distribution using all data from the TRF at negative delays from −40 ms to 0 ms and calculated empirical *p*-values for each time lag at the population level.

An ANOVA was conducted to assess differences in speech comprehension across the different audiovisual stimulus conditions. Following significant ANOVA results, pairwise comparisons between conditions were performed using paired *t*-tests.

An ANOVA was also used to test for differences in the TRF peaks between the four audiovisual conditions. Additionally, repeated-measures correlations were employed to assess the putative relationship between the neural data and each participant’s speech comprehension score.

The neural response was also analysed for lateralization. Therefore, the signal strength at the population-level peak latency was extracted for each voxel separately. The data were normally distributed according to the Shapiro-Wilk test. Consequently, the signal strengths between the source points in the left hemi-sphere and in the right hemisphere were compared with each other through a *t*-test followed by Bonferroni correction for multiple comparisons.

## Results

### Speech Comprehension

We determined the average comprehension scores of the participants on the population level, but considered them separately for each of the four types of audiovisual stimuli. We reported the resulting speech com-prehension scores already in our previous work on this dataset [Riegel et al., 2026]. A repeated-measures ANOVA across all audiovisual conditions revealed a significant main effect of condition on speech compre-hension (*p <* 0.001).

When participants saw a still image of the speaker while listening to the sentence, they understood, on average, 66% of the words correctly (Figure 2). Their understanding increased significantly (*p <* 0.001) to an average of 78% when they watched an avatar of the speaker while listening to the spoken sentences. The same increase (*p <* 0.001) was observed between the still image and the degraded video, in which only strong edges in the speaker’s face were visible. This condition was designed to serve as a control version of the avatar. This worked well, as the participants correctly understood 76% of the words, which did not differ significantly from the avatar condition. Comprehension increased significantly again (*p <* 0.001) when watching the natural video, with which the participants correctly understood, on average, 84% of the words.

**Figure 1:**
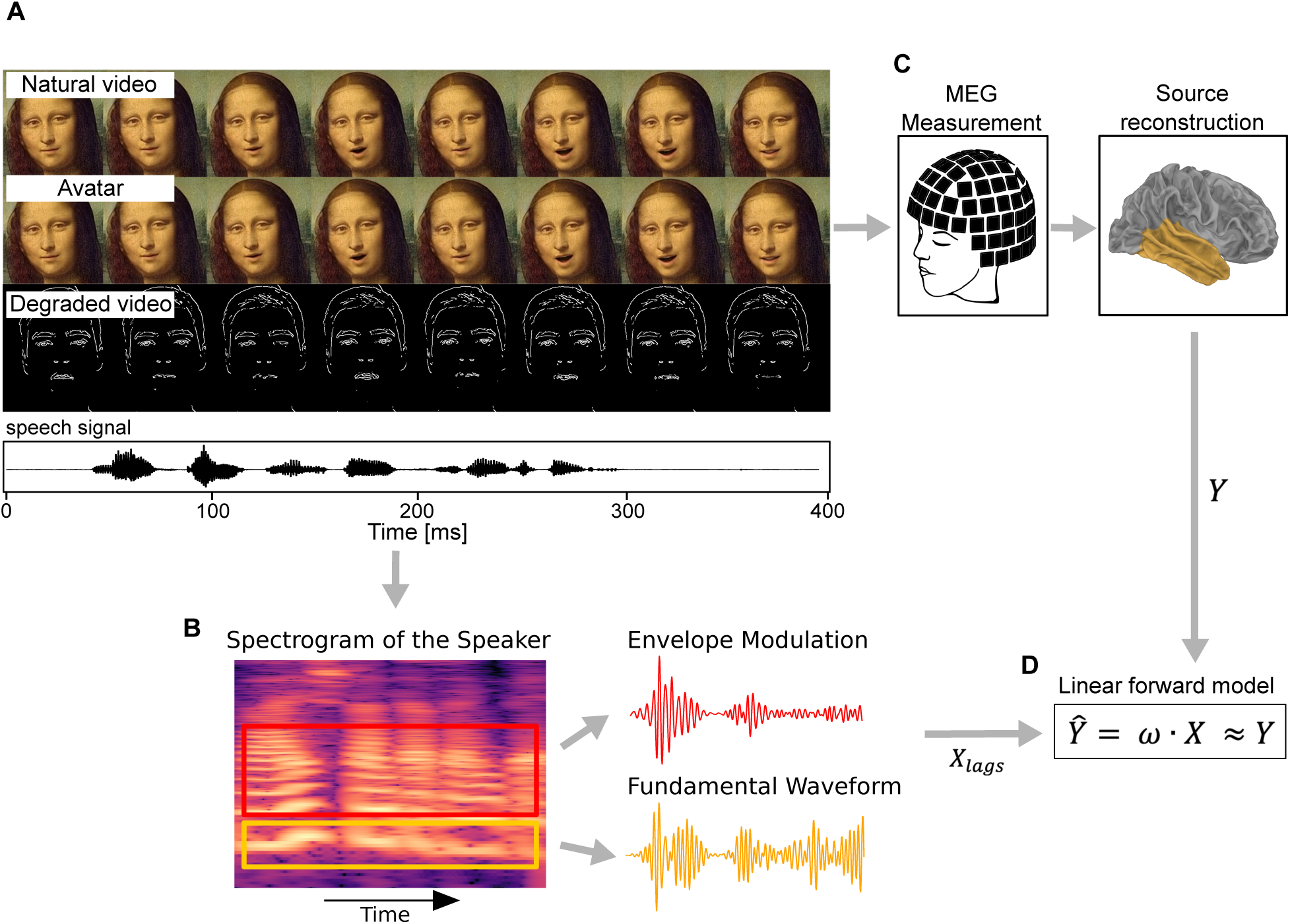
Overview of the experimental procedure and audiovisual stimuli. (A)The images of the real speaker and his avatar are replaced in this Preprint by a speaking avatar of Mona Lisa. Three types of audiovisual stimuli were generated for this study: the speaker’s natural video, an avatar generated with DNNs from a still image and an audio file, and a degraded version of the natural video obtained by applying an edge detection algorithm. Below the frames, the corresponding speech signal is shown. (B) Two features (*X_lags_*) are extracted from the spectrogram of the speech signal: a representation of the fundamental waveform that oscillates at the fundamental frequency (yellow), and the envelope modulation of its higher harmonics (red). (C) MEG measurements were conducted while participants watched the stimuli. The MEG data were source-reconstructed onto the auditory cortex (*Y* ). (D) The speech-FFR is determined by estimating the source-reconstructed MEG data (*Y*^^^ ) from the two speech features (*X_lags_*) using a linear forward model. The magnitude of the resulting coefficients (*ω*) over time yield the the Temporal Response Functions (TRFs).

**Figure 2:**
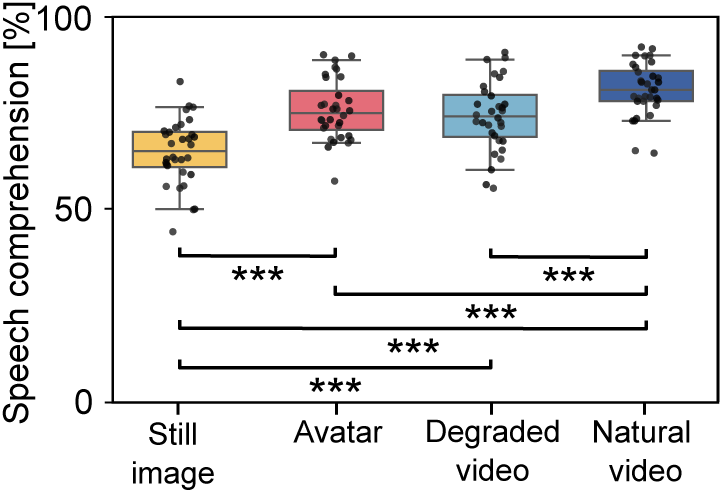
Speech comprehension scores for each audiovisual condition across participants. Comprehension was lowest for the audio-only condition, higher for the avatar and degraded video, and most words per sentence were understood for the natural video. All increases in comprehension were strongly significant (*p <* 0.001) except between the avatar and its control condition, the degraded video.

### Speech-FFR

We determined the speech-FFR by relating two speech features to the MEG data. The first feature, the fundamental waveform, captured neural activity phase-locked to the fundamental frequency of the speech. The second feature, the envelope modulation of the higher harmonics, captured the power fluctuations in the higher harmonics at the fundamental frequency and the related neural response.

We found that both features produced significant peaks in their temporal response functions (TRFs, Fig-ure 3). The TRFs exhibited comparable behaviour across the four types of audiovisual conditions. Neural responses to the fundamental waveform peaked between 30 ms and 36 ms (Figure 3*A*), whereas responses to the envelope modulation occurred slightly earlier between 26-30 ms (Figure 3*B*).

**Figure 3:**
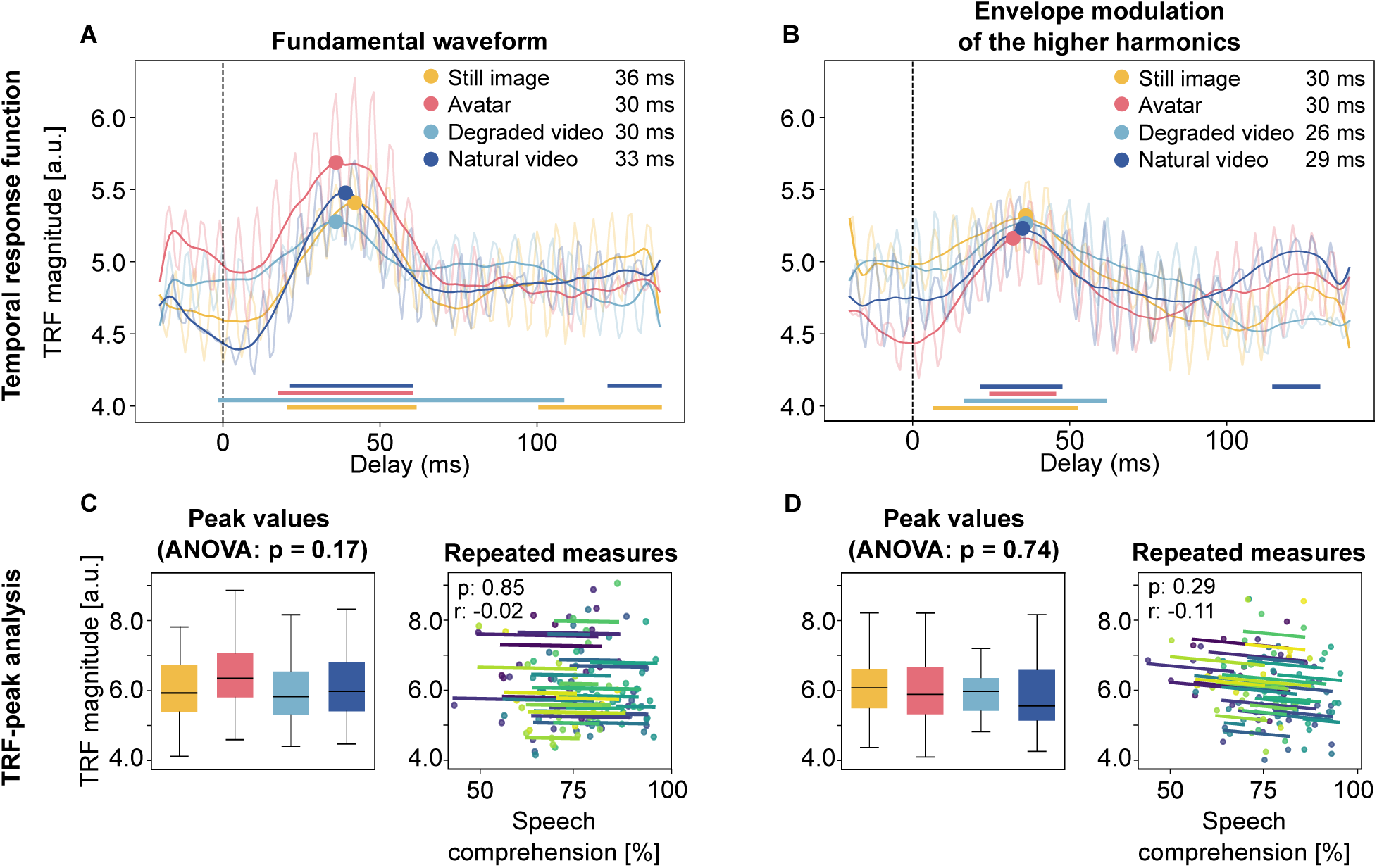
Temporal response functions (TRFs) in the four different audiovisual conditions. **A, B**, TRFs for the fundamental waveform (**A**) and for the envelope modulation of the higher harmonics (**B**) for all four audiovisual conditions. The transparent lines show the amplitude of the TRFs, and the opaque lines are the low-pass filtered version of the amplitudes. The dots in the plots represent the peaks of the low-pass filtered TRFs. The timing of the peaks is given in the legend. The bars at the bottom indicate at which time lags the TRFs are significant. **C, D**, The TRF-peak analysis for both the fundamental waveform (**C**) and the envelope modulation of the higher harmonics (**D**). The left box plot in both (**C**) and (**D**) shows the magnitudes of the TRF peaks across participants for each audiovisual condition. The right plot shows the repeated-measures correlation between the magnitude of the TRF peaks and the behavioral scores across the four conditions. Each color represents one participant.

To assess whether audiovisual integration was apparent in the speech-FFR, we compared the response strength across the different audiovisual conditions. Therefore, ANOVA was used to test for significance in the distribution of the peak magnitudes of the individual participants across all conditions. Neither for the fundamental waveform (Figure 3*C*, *left*) nor for the envelope modulation (Figure 3*D*, *left*) did these yield significant results (fundamental waveform: *p* = 0.17; envelope modulation: *p* = 0.74).

Although a direct comparison of the magnitude of the speech-FFR across the four conditions did not produce significant results, we further examined whether the trend in neural signal strength correlated with the speech comprehension scores. We therefore computed repeated measures correlations. We again did not obtain a statistically significant result (fundamental waveform: *p* = 0.85, Figure 3*C*, *right*; envelope modulation: *p* = 0.29, Figure 3*D*, *right*).

### Lateralization

In addition to analysing the source-reconstructed MEG data averaged across the whole auditory cortex, we also investigated lateralization between the left hemisphere (LH) and the right hemisphere (RH). We therefore compared the neural activity at the population-level peak latencies, averaged over all participants. We found a stronger neural response in the right hemisphere, both to the fundamental frequency and to the envelope modulation of the higher harmonics (Figure 4).

**Figure 4:**
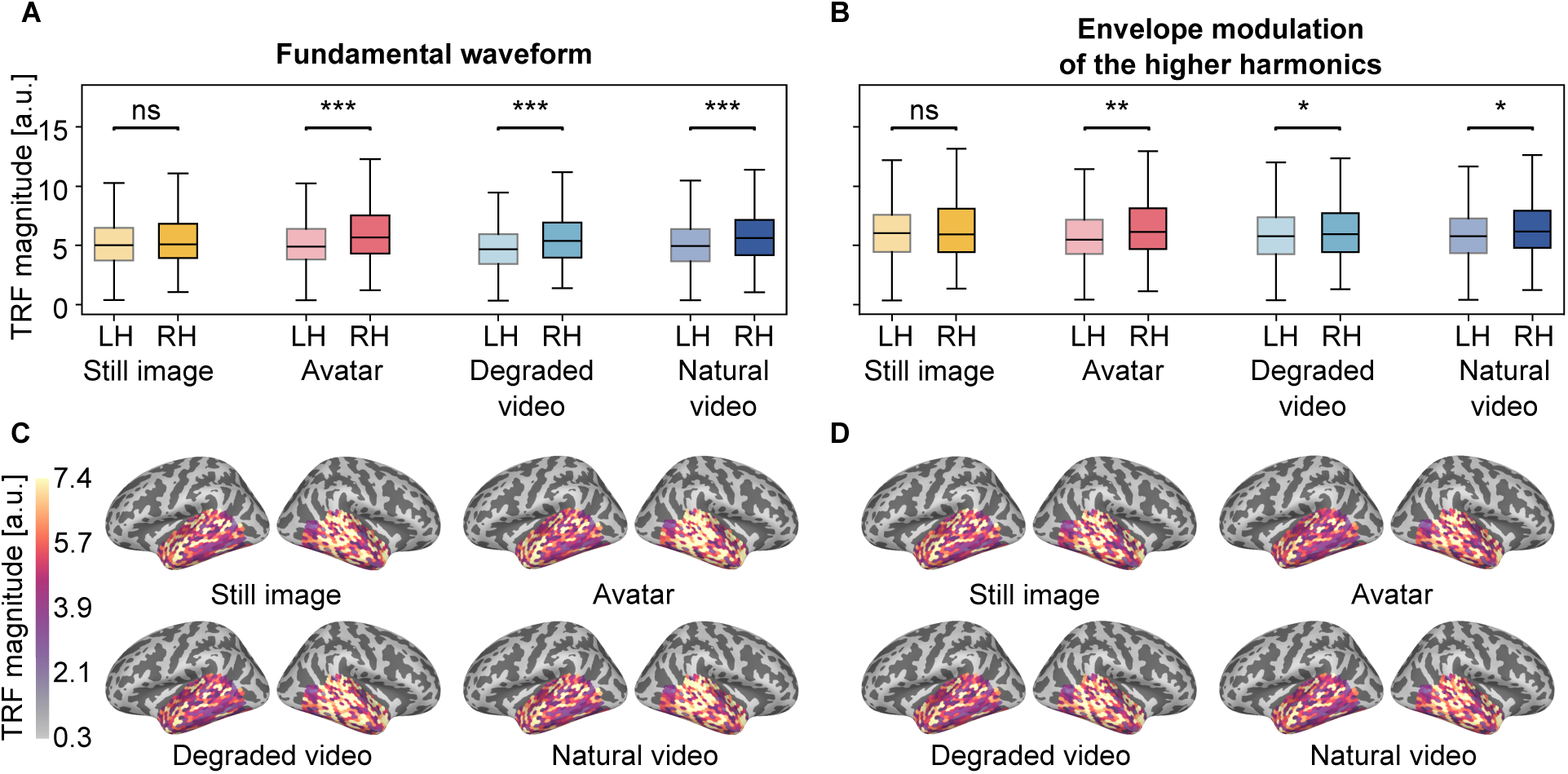
Lateralization of the speech-FFR in the four audiovisual conditions at the lag of the population TRF peak. **A, B**, Per condition, the lighter, left box shows the neural response to the predictor variable at the population level in the left hemisphere across voxels. The darker box on the right represents the corresponding activation in the right hemisphere. **A**, The response to the fundamental waveform in the still image condition shows a non-significant tendency of right lateralization. All conditions with video show a strong, significant right lateralization. **B**, The neural response to the envelope modulation of the higher harmonics is not significantly lateralized for the still image. In all conditions containing a video, the neural response exhibits significant right lateralization. **C, D**, Neural activation in the auditory cortex in lateral and medial views is presented in each condition. Both the response to the fundamental waveform (**C**) as well as the envelope modulation of the higher harmonics (**D**) appear right lateralized on visual inspection.

Although the difference is visible as a trend across all four audiovisual conditions, it is significant only in the three conditions that include visual-speech information (Bonferroni-corrected). The neural response to the fundamental waveform is significantly right lateralized (*p <* 0.001) in these three conditions (Figure 4*A*) (audio-only: *p* = 1.0).

The lateralization of the response to the envelope modulation is also significant in all AV conditions (avatar: *p* = 0.006, degraded: *p* = 0.01, and natural: *p* = 0.04). In contrast, the lateralization of the audio-only condition (*p* = 1.0) is not significant.

To check whether the strength of lateralization per participant was significantly correlated with participants’ speech comprehension, we calculated a lateralization score *L* for each participant as 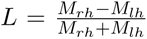, with *M_rh_* denoting the magnitude of the TRF peak in the right hemisphere and *M_lh_* that in the left hemisphere. We then computed the repeated measures correlation between this lateralization score *L* and the participants’ speech comprehension score in each of the four audiovisual conditions. Again, this did not yield a significant effect (fundamental waveform: *p* = 0.75; envelope modulation: *p* = 0.21).

## Discussion

We investigated whether the cortical contribution to the speech-FFR is influenced by seeing a speaker’s face. We not only presented the speaker’s natural video but also a degraded version of the video and an avatar that resembles the speaker. Background noise was employed to reduce speech comprehension and to make the audiovisual integration apparent. This strategy worked as desired: the participants understood significantly more of the spoken words with the visual signals than without. Moreover, speech comprehension for the natural videos exceeded that for the avatar and the degraded video.

Several recent studies investigated how well avatars can support speech understanding[Munhall et al., 2004, Varano et al., 2022, Shan et al., 2022]. The findings reported here are in line with these previous studies, showing that such avatars yield speech comprehension scores that lie between those for the audio-only condition and for the natural video. Although many participants did not notice a difference between the avatar and the natural video (undocumented comments by the participants), the behavioral difference in speech comprehension between the avatar and the natural video was highly significant. This is probably not only due to the lips of the avatar not articulating perfectly, but also due to the timing between lip movement and speech not being fully accurate. In contrast, the degraded video resulted in the same level of speech comprehension as the avatar. Although the degraded video had, by construction, perfect temporal alignment between lip movement and audio signal, it lacked a realistic appearance.

Regarding neural responses, we found significant peaks around the expected latency of 30 ms, as identified in previous studies [Schüller et al., 2024, 2025]. This timing confirmed that we could reliably detect the cortical contribution to the speech-FFR, despite the comparably high background noise level (SNR of - 4 dB)[Schüller et al., 2024]. However, when comparing the peak magnitudes across the different audiovisual conditions, no statistically significant differences were observed.

To the best of our knowledge, ours is the first study to investigate the cortical contribution to speech-FFR elicited by audiovisual speech, as well as the first to measure this neural response when elicited by an artificially-generated avatar. Existing evidence for audiovisual influences on high-frequency auditory processing largely comes from studies using simplified, speech-like stimuli. In particular, Musacchia et al. [2006] and Musaccia et al. [2007] examined the effect of audiovisual presentation of the syllable / da / on subcortical responses measured with EEG and reported clear suppression of the speech-FFR through the visual signal at very early latencies. At the cortical level, an ECoG study by Karthik et al. [2021] investigated responses to the same type of syllabic stimulus and observed larger neural activity in the high-gamma band in the presence of the visual signal. However, these effects were restricted to a very narrow temporal window closely aligned with stimulus onset and were spatially very limited.

Importantly, both Musacchia et al. [2006] and Karthik et al. [2021] employed audio-only stimuli that did not include a realistic visual depiction of the speaker’s face. In contrast, our paradigm presents a static image of the speaker in the audio-only condition, which may reduce the visual contrast between conditions with and without video, and thereby limit the magnitude of the observable audiovisual effects. Furthermore, our experimental setup more closely resembles real-world listening situations, as it involves continuous speech and relatively strong background babble noise. Given that high-gamma audiovisual effects were only marginally detectable in very precise areas within the STG [Karthik et al., 2021] even under highly controlled conditions measured with electrocorticography, it is plausible that such effects are not sufficiently robust to be observed in our more challenging and naturalistic MEG paradigm. In particular, we note that the relatively high level of background noise that we employed reduces the magnitude of the speech-FFR compared to studies with less background noise, which may make putative audiovisual effects harder to detect. Notably, Karthik et al. [2021] reported more robust audiovisual modulations in lower frequency bands, effects that we have also reported separately for our data [Riegel et al., 2026].

In addition to analyzing the magnitude of the speech-FFR, we also examined its hemispheric lateralization. Across all conditions, responses showed a clear right-hemispheric bias. Previous audio-only studies have similarly reported right-lateralized cortical responses to the fundamental waveform and the envelope modulation of the higher harmonics of continuous speech [Schüller et al., 2023]. Interestingly, in our study, lateralization was significant only for stimuli with visual information. We would expect the lateralization trend of our audio-only stimuli to also reach significance in an experimental paradigm with lower background noise. Our finding of a somewhat stronger lateralization for the stimuli with visual input could hint at a minor effect of audiovisual integration in the high-gamma band.

In summary, we examined the cortical contribution to the speech-FFR elicited by audiovisual speech. The magnitude of this response did not depend on the type of visual stimulus, and was also identical when participants watched an artificially-created avatar. Despite substantial variation in the speech-in-noise comprehension scores between the participants and the audiovisual conditions, we found that these could not be explained by the speech-FFR. Our results suggest that audiovisual integration occurs at later cortical stages. Future investigations may further clarify the latencies at which audiovisual integration regarding speech stimuli appears, and whether pitch processing may be involved.

## Data Accessibility

The data used in this study are available on Zenodo: https://zenodo.org/records/18258659

## Acknowledgments

This project was supported by the German Science Foundation (DFG, project number 523344822).

## Author contributions

JR performed research, collected data, analysed the data, interpreted the results, and wrote the paper. AS collected the data. AW, SZ, DK, and TR designed research, interpreted the results, and edited the paper.

